# Till evolution do us part: The diversity of symbiotic associations across populations of *Philaenus* spittlebugs

**DOI:** 10.1101/2022.12.05.519182

**Authors:** Michał Kolasa, Łukasz Kajtoch, Anna Michalik, Anna Maryańska-Nadachowska, Piotr Łukasik

**Affiliations:** Institute of Environmental Sciences, Faculty of Biology, Jagiellonian University, Krakow, Poland; Institute of Systematics and Evolution of Animals, Polish Academy of Sciences, Krakow, Poland; Department of Developmental Biology and Morphology of Invertebrates, Institute of Zoology and Biomedical Research, Faculty of Biology, Jagiellonian University, Krakow, Poland

## Abstract

Symbiotic bacteria have played crucial roles in the evolution of sap-feeding insects and can strongly affect host function. However, their diversity and distribution within species are not well understood; we don’t know to what extent environmental factors or associations with other species may affect microbial community profiles. Here, we sequenced host and bacterial marker gene amplicons to survey the bacterial community composition across multiple populations of *Philaenus* spittlebugs.

Host mitochondrial sequence data confirmed morphology-based identification of 6 species and revealed two divergent clades of *Philaenus spumarius*. All of them hosted the primary symbiont *Sulcia* that was almost always accompanied by *Sodalis*. Interestingly, populations and individuals often differed in the presence of *Sodalis* sequence variants, suggestive of intra-genome 16S rRNA variant polymorphism combined with rapid genome evolution and/or recent additional infections or replacements of the co-primary symbiont. The prevalence of facultative endosymbionts, including *Wolbachia, Rickettsia*, and *Spiroplasma*, varied among populations.

Notably, COI amplicon data also showed that nearly a quarter of *P. spumarius* were infected by parasitoid flies (*Verralia aucta*). One of the *Wolbachia* OTUs was exclusively present in *Verralia*-parasitized specimens, suggestive of parasitoids as their source and highlighting the utility of host gene amplicon sequencing in microbiome studies.

## Introduction

Symbiosis with microorganisms has been a critical force driving eukaryotic evolution. Through their effects, ranging from the biosynthesis of nutrients, through defense against pathogens, to the manipulation of reproduction (Moran *et al*., 2008; Feldhaar, 2011), symbiotic microorganisms have repeatedly enabled the emergence of significant insect clades feeding on specialized foods, as well as allowing for more dynamic responses to spatial heterogeneity, natural enemy pressure, and changing environmental conditions (Jaenike *et al*., 2010; Smith *et al*., 2021). Although our knowledge expanded significantly during the last two decades, we are still far from understanding the relationship dynamics between symbionts and their hosts. Fortunately, novel methodological and conceptual developments help consolidate our understanding of these interactions.

When considering symbiotic interactions, it is crucial to consider their nature and stability. The recent classification of symbioses into closed, open, and mixed (Perreau and Moran, 2021) has created a helpful framework for such comparisons. Closed symbioses, usually characterized by their ancient origin, genome reduction of the symbiont, and its strict vertical transmission through host generations (Bennett and Moran, 2015), include heritable nutritional symbionts of sap-feeding hemipterans that provide them with essential amino acids and other nutrients deficient in their unbalanced diet (Douglas, 2009, 2016; McCutcheon *et al*., 2009). Auchenorrhyncha, one of the major hemipteran clades that date back some 300 million years and include planthoppers, leafhoppers, treehoppers, spittlebugs, and cicadas, were ancestrally infected by such symbiont, a Bacteroidetes, *Candidatus* Sulcia muelleri (further referred to as *Sulcia*). In major Auchenorrhyncha clades, *Sulcia* was ancestrally accompanied by co-symbionts such as a betaproteobacterium *Zinderia* in spittlebugs (McCutcheon and Moran, 2010), or alphaproteobacterium *Hodgkinia* in cicadas (McCutcheon *et al*., 2009). However, these ancient microbes, co-diversifying with hosts, have been repeatedly replaced by other microbes. For example, in many clades of cicadas, *Hodgkinia* was replaced by *Ophiocordyceps* fungi which took over its nutritional roles, while *Sulcia* remained without major changes (Matsuura *et al*., 2018). Similarly, in the Philaenini tribe of spittlebugs, *Zinderia* was not found, and all studied individuals hosted *Sodalis*-allied bacteria instead. Genomic data revealed that in *Philaenus spumarius, Sodalis* has taken over the functions of *Zinderia* in the common ancestor of *P. spumarius*, and, apparently, other Philaenini (Koga and Moran, 2014).

Another functional category, mixed symbioses, is not fixed on evolutionary time scales. In addition to relatively reliable maternal transmission, facultative endosymbionts are known for infecting new hosts and spreading in populations rapidly, even on the continental scale (Himler *et al*., 2011; Kriesner *et al*., 2013; Turelli *et al*., 2018). While historically regarded as reproductive parasites (Hurst and Jiggins, 2000), widespread symbionts such as *Wolbachia, Rickettsia, Spiroplasma*, and *Cardinium* can influence host adaptations in multiple ways. *Wolbachia*, the best known and the most prevalent facultative endosymbiont that is estimated to infect ca. 40% of all arthropods (Zug and Hammerstein, 2012), can upregulate hosts’ immune genes (Hedges *et al*., 2008; Kambris *et al*., 2010), protect them from natural enemies (Gerardo and Parker, 2014), influence thermal preference (Truitt *et al*., 2019), and susceptibility/resistance to insecticides (Liu and Guo, 2019) or even supply the host with nutrients (Hosokawa *et al*., 2010; Ju *et al*., 2020). Although less studied and not as broadly distributed, other facultative endosymbionts can have a similar range of effects (e.g., Chiel *et al*., 2009; Brumin *et al*., 2011; Łukasik *et al*., 2013; Xie *et al*., 2014; Liu and Guo, 2019).

Insects can also host diverse microbes in their digestive tracts and on body surfaces. Many of them belong to the third “open symbiosis” category, generally not forming long-term associations with hosts. Gut microbiota is not generally thought to play significant roles in the biology of sap-feeding insects, even though this has not been comprehensively investigated (Buchner, 1965; Engel and Moran, 2013; Luo *et al*., 2021).

The recent development of high-throughput sequencing has led to the blooming of microbiome studies, substantially expanding our understanding of the diversity of insect symbioses. However, while increasingly understood, the numerous methodological caveats associated with tools such as 16S rRNA amplicon sequencing are not often systematically addressed (Knight *et al*., 2018). For example, wild insect microbiome studies tend to neglect issues such as molecular confirmation of the host identity or the presence of parasites and parasitoids. A completely different problem is reagent- and cross-contamination, which can be a major problem, especially in samples with low bacterial load (Salter *et al*., 2014), and is hard to tackle without negative controls for different laboratory steps. However, these issues can be resolved. We can largely avoid these issues by amplifying host marker genes alongside those of microbes, using negative controls and well-thought bioinformatic pipelines (Knight *et al*., 2018).

The primary focus of the present study is the characterization of the diversity and distribution of the microbiota of the meadow spittlebug *P. spumarius* (Hemiptera: Aphrophoridae). This polyphagous xylem-feeding insect of Palearctic origin was recently introduced in North America, the Azores islands, Hawaii, and New Zealand (Rodrigues *et al*., 2014). As a vector of *Xylella fastidiosa*, a recently emerged and dangerous pathogen of crops including olives, grapevines, citrus trees, and coffee, *P. spumarius* has become an economically important species, especially in Europe (Saponari *et al*., 2014; Godefroid *et al*., 2021). Its microbiota, critical to spittlebugs’ nutrition and likely affecting other functions, perhaps including interactions with vectored plant pathogens, have thus attracted researchers’ attention.

Members of the Philaenini tribe of spittlebugs were found to be universally infected by *Sulcia* and *Sodalis* symbionts (Koga *et al*., 2013) which together produce ten essential amino acids deficient in the xylem sap diet (Koga and Moran, 2014). Koga and Moran (2014) postulated the establishment of *Sodalis* as a nutritional symbiont in the ancestor of the whole *Philaenini* tribe. However, they comprehensively characterized only a single *P. spumarius* population, and their work was limited to a few samples. On the other hand, much attention has been paid recently to understanding the distribution and dynamics of the facultative symbionts such as *Wolbachia, Rickettsia*, and *Cardinium* across different geographic populations of the *Philaenus* group, especially in the Mediterranean region (Lis *et al*., 2015; Kapantaidaki *et al*., 2021; Formisano *et al*., 2022). However, the picture of microbiota across the *Philaenus* diversity, including the stability of the association with nutritional symbionts, the distribution of facultative symbionts, or the presence of other bacterial categories, is far from complete.

Here we aim to characterize the diversity and distribution of symbiotic bacteria in six species belonging to the genus *Philaenus*, with a strong emphasis on different populations of *P. spumarius*. We use insect and symbiont marker gene amplicon sequencing to address questions about the stability of obligatory nutritional “closed” symbioses and the diversity and distribution of facultative endosymbionts across species and populations. We discuss the symbiont diversity and distribution patterns, with emphasis on the roles of parasitoids and the process of symbiont replacement.

## Materials and Methods

### Sampling

*Philaenus* species were collected during field trips across Europe, the Middle- and the Far East between 2000 and 2011 (Fig. 1, Suppl. Tab. 1). Specimens were preserved in 96% ethanol and stored at −20°C until processing. Morphology-based identifications by expert taxonomists revealed eight species: *P. spumarius* (22 populations / 72 specimens), *P. signatus* (3 populations / 7 specimens), and a single population of other species: *P. tesselatus* (3 specimens), *P. arslani* (2 specimens), *P. italosignus* (2 specimens), *P. loukasi* (2 specimens), *P. maghresignus* (1 specimen), and *P. tarifa* (1 specimen). In total, 90 specimens from eight species were chosen for amplicon-based characterization. However, as explained later, molecular barcodes did not support some of these identifications, which we decided to use as the primary means to determine species.

**Figure 1.**
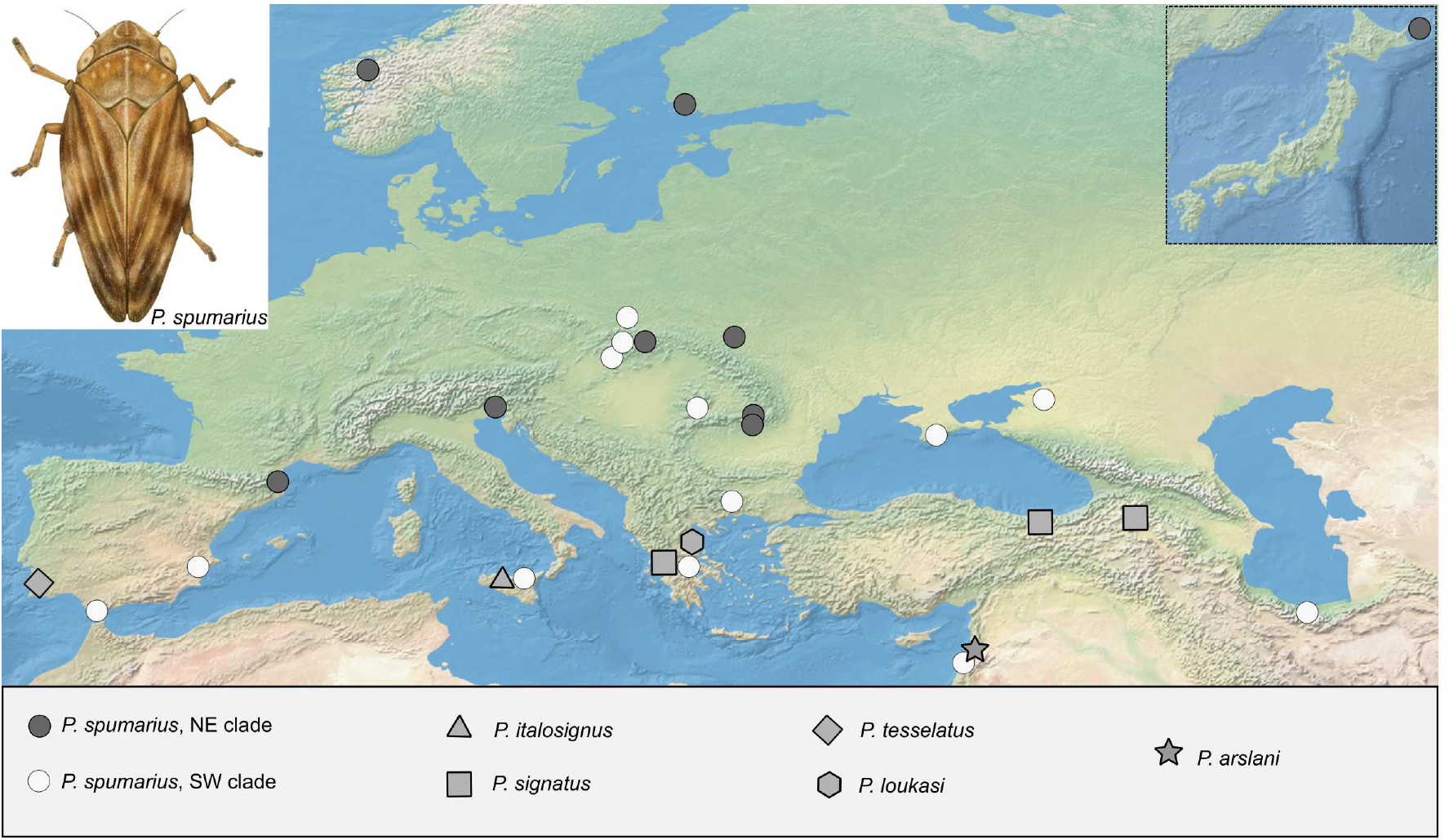
The sampling sites of specimens from the *Philaenus* group classified into species based on mitochondrial sequences. Circles - *P. spumarius*, triangle - *P. italosignus*, rectangle - *P. signatus*, diamond - *P. tesselatus*, hexagon - *P. loukasi*, five-pointed star - *P. arslani*. The box in the upper right corner shows the sampling site of *P. spumarius* on Kunashir Island, Japan. Colors indicate mitochondrial clades of P. spumarius: white - NE, dark grey - SW.

### Host diversity and microbiome screen

#### Library preparation and sequencing

We characterized the selected insects using a custom two-marker-gene amplicon sequencing approach. We simultaneously targeted the insect mitochondrial cytochrome I oxidase (COI) gene, allowing for the confirmation of the host identity and the V4 hypervariable region of the bacterial 16S rRNA gene, providing a picture of its microbiota.

DNA was extracted from whole insects using the Nucleospin Tissue kit (Macherey-Nagel), following the manufacturer’s instructions. The extraction was conducted in two batches: some samples were processed back in 2009 and others in December 2019 (Suppl. Tab. 1). Samples and negative extraction controls for both batches were used for amplicon library preparation following a modified two-step PCR library preparation approach as outlined by Glenn (2011) (method 4). In the first step, two marker regions of interest were amplified using template-specific primers 515F/806R (Apprill *et al*., 2015; Parada *et al*., 2016) and COIBF3/COIBR2 (Elbrecht *et al*., 2019) with Illumina adapter tails. The PCR products were purified using SPRI magnetic beads and used as templates for the second indexing PCR reaction. Pooled libraries were sequenced on an Illumina MiSeq v3 lane (2 × 300 bp reads) at the Institute of Environmental Sciences of Jagiellonian University. The primer sequences and detailed protocols for amplicon library preparation are provided in Supplementary Table 2.

#### Analyses of amplicon sequencing data

We processed 16S rRNA and COI amplicon data using a custom pipeline based on USEARCH/VSEARCH (described in detail and available at https://github.com/Symbiosis-JU/Philaenus-Microbiota-Project). Initially, all amplicon datasets were split into bins corresponding to the two target genes based on primer sequences. Using PEAR, we assembled quality-filtered forward and reverse reads for both bins into contigs (Zhang *et al*., 2014). Next, contigs were de-replicated (Rognes *et al*., 2016) and denoised (Edgar, 2016); this was done separately for every library to avoid losing information about rare genotypes that could happen during the denoising of the whole sequence set at once (Prodan *et al*., 2020). The sequences were then screened for chimeras using USEARCH and then classified by taxonomy using the SINTAX algorithm and customized databases: SILVA for bacterial 16S rRNA (version 138 SSU) and MIDORI (version GB 239) for host COI, which were screened for errors. Finally, the sequences were clustered at a 97% identity level using the UPARSE-OTU algorithm implemented in USEARCH. The resulting Operational Taxonomic Unit (OTU) tables were then used for a series of custom steps. Bacterial 16S rRNA gene data were screened for putative DNA extraction and PCR reagent contaminants using a set of negative controls (blanks) as a reference.COI data were screened for bacterial and parasite reads using custom references. Specimens with a low number of COI or 16S V4 reads after decontamination (not included in the counts above) were excluded from further analysis.

The Minimum Spanning Network for COI haplotypes of *Philaenus spumarius* was generated using PopArt (Leigh and Bryant, 2015).

### Microscopic analyses of symbiont morphology and localization

#### Light (LM) and transmission electron (TEM) microscopy

Abdomens of *P. spumarius* females collected in Krakow (Poland) were fixed and stored in 2.5% glutaraldehyde solution in 0.1 M phosphate buffer (pH 7.4) at 4°C for three months and, after this time, washed with the same buffer with the addition of sucrose (5.8%). Next, samples were postfixed in a 1% solution of osmium tetroxide, dehydrated in ethanol and acetone series, and embedded in epoxy resin Epon 812 (SERVA, Heidelberg, Germany). The resin blocks were cut into serial, semithin sections for histological analyses, stained in 1% methylene blue in 1% borax, and observed under the Nikon Eclipse 80i light microscope. Ultrathin sections for ultrastructural analyses were contrasted with lead citrate and uranyl acetate and observed under the JEOL JEM 2100 electron transmission microscope.

#### Fluorescence microscopy (FM)

For fluorescence *in situ* hybridization (FISH), ethanol-preserved specimens of *P. spumarius* (from Spain) were rehydrated, postfixed in 4% paraformaldehyde for two hours, and then dehydrated again through incubation in increasing concentration of ethanol and acetone. The samples were then embedded in Technovit 8100 resin (Kulzer, Wehrheim, Germany) and cut into semithin sections. Hybridization using *Sulcia-* and *Sodalis*-targeting probes labeled with Cy3 and Cy5 fluorochromes (sequences in Suppl. Table 2) was performed overnight at room temperature. Slides were washed three times in PBS solution, dried, covered with ProLong Gold Antifade Reagent (Life Technologies), and examined using a confocal laser scanning microscope Zeiss Axio Observer LSM 710.

## Results

### COI amplicon data permit validation of species identity and reveal parasitoid infections

After filtration, the total COI reads taxonomically assigned to Auchenorrhyncha was 570,638, or 6,341 per sample on average. We identified 34 amplicon sequence variants (ASVs) that clustered to 6 operational taxonomic units (OTUs) with a 97% identity cutoff. The vast majority of examined specimens were characterized by a dominant COI genotype confidently classified as representing the genus *Philaenus*. Out of 90 libraries, in 71, the dominant sequence variant from that OTU represented over 95% of total COI reads; in further 8, it represented 90 - 95% of reads, and >70% in the remaining 11. Widespread secondary ASVs with the same species-level assignments were found in *P. spumarius* (2 ASVs) or *P. signatus* (one ASV). They were much less abundant and their relative abundances were similar in different specimens, suggestive of them representing nuclear pseudogenes (Dong et al., 2021) (Fig. 2, Table 3).

**Figure 2.**
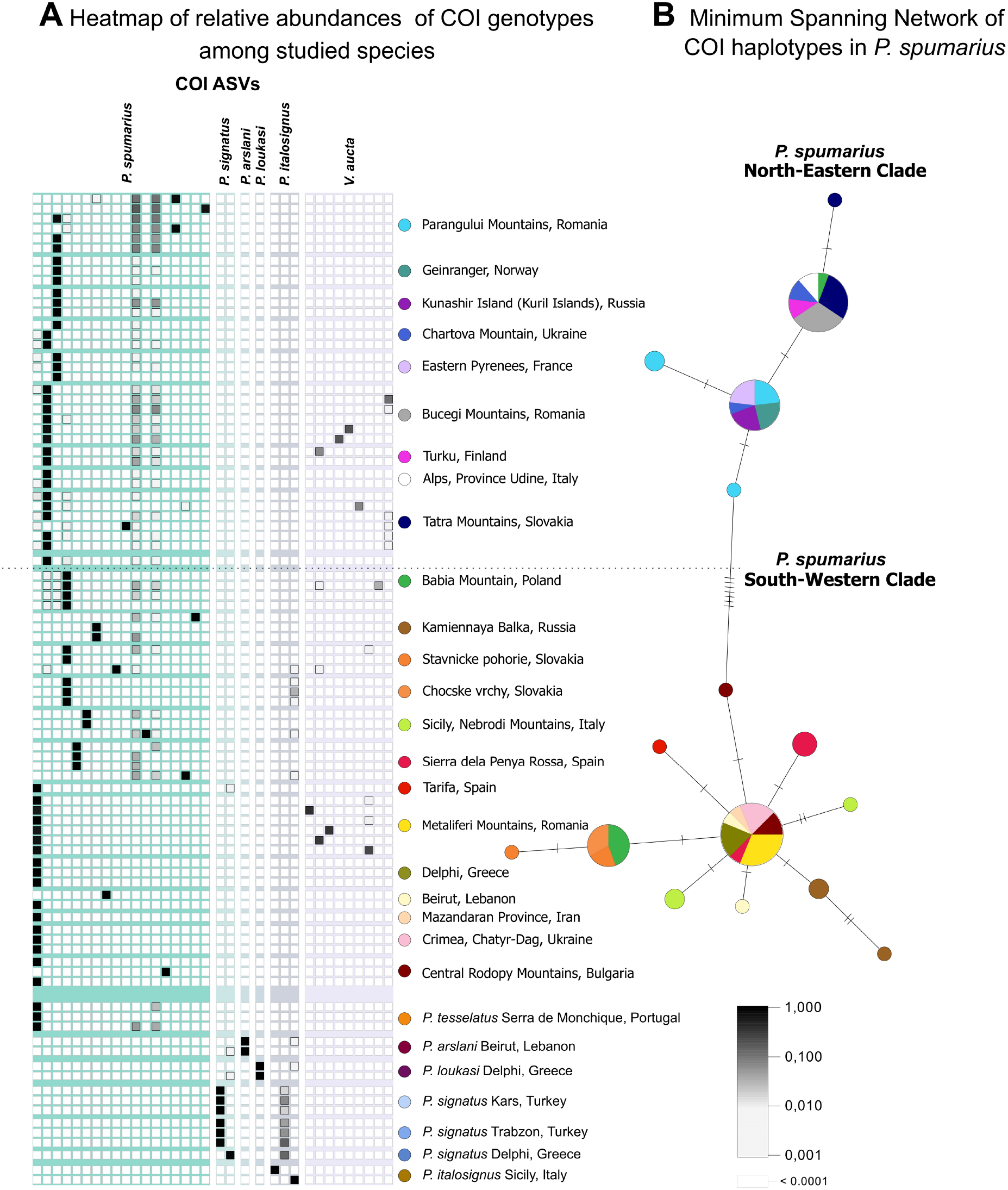
**A** - Distribution of Cytochrome Oxidase subunit I (COI) genotypes (ASVs) among studies species. **B** - Minimum Spanning Network of COI haplotypes of *P. spumarius*. The dashed line indicates the separation between NE clade and SW clade of *P. spumarius*. Colored circles indicate populations from which samples were collected and congruent with the haplotype network colors.

Interestingly, in 19 samples of *P. spumarius*, in addition to the spittlebug signal, we found one of 9 ASVs belonging to a single 97% OTU identified as representing the parasitoid fly *Verralia aucta* (Diptera: Pipunculidae). In 9 of those samples, parasitoid reads exceeded 1% of the total.

Analyzes of COI marker gene sequences for the experimental specimens confirmed their morphology-based identifications in most cases. We decided to retain the morphology-based delimitation of *P. tesselatus* and *P. spumarius*, previously shown to share the same COI haplotype (Maryańska-Nadachowska *et al*., 2012) (Figure 2A). However, specimens labeled as *P. maghresignus* and *P. tarifa* were indistinguishable from *P. spumarius* based on their COI gene fragment (despite previously being shown to be distinct); therefore, we classified them as *P. spumarius*. We later verified that their microbiota did not depart from those of *P. spumarius* from the same or nearby sites.

The final dataset contains 74 specimens assigned as *P. spumarius*, seven as *P. signatus*, three as *P. tesselatus*, and two specimens per species as *P. arslani, P. loukasi*, and *P. italosignus*. The haplotype networks revealed two geographically disjunct clades of *P. spumarius* (Fig. 2B) with hybridization zones in the Carpathian arc, in agreement with previous studies (Lis *et al*., 2014).

In a subset of specimens, including all individuals of *P. signatus*, we also discovered COI ASVs corresponding to other species of *Philaenus* than the dominant genotype - specifically, to *P. italosignus*. In all but one case, their cumulative relative abundance did not exceed 1%. In *P. signatus*, the sequence is different from those present in studied *P. italosignus*, suggestive of being a NUMT (Song *et al*., 2008), but in other cases, it matches the barcode of one *P. itolosignus* specimen. We verified that there was no apparent cross-contamination in bacterial 16S rRNA amplicon data generated in the same PCR reactions and conclude that the observed cross-specific signal is likely a biological phenomenon (Fig. 2A).

### Clear patterns in the distribution of bacterial genera

The comparison of 16S rRNA gene V4 region data for spittlebugs against PCR and extraction blanks showed that, on average, 99.83% of reads in the library represented actual insect-associated microbes rather than contaminants (Suppl. Table 5). The highest contamination level observed in the sample was 3.7%. Reads classified as representing reagents or laboratory contaminants were removed.

The total number of 16S rRNA reads after all analysis steps were 1,488,428, or 16,538 per sample on average (range 1,098 - 47,801). Within these data, we identified 217 ASVs (zOTUs) clustered to 89 OTUs with a 97% identity cutoff. The dominant microbial OTUs represented taxa previously reported from *P. spumarius* or other insects. All studied *Philaenus* individuals hosted *Sulcia*, and 88 out of 90 hosted *Sodalis*. Together, these obligate symbionts comprised 78.9% of all bacterial reads in a library on average. Other bacteria known as insect-associated were present in some specimens. We detected *Wolbachia* in 46.6%, *Rickettsia* in 15.5%, *Spiroplasma* in 6.6%, and *Pectobacterium* in 15.5% of individuals. Together, these six clades comprised 96.5% of reads in a library on average. Other, less prevalent and abundant microbes, including the genus *Pseudomonas* and the families Oxalobacteraceae and Rhizobiaceae, represent taxa reported from a variety of environments, including insects.

*Pectobacterium*, an enterobacterial symbiont known to colonize various invertebrates as a nutritional heritable mutualist (Martinson *et al*., 2020), was found in a small fraction of *P. spumarius* individuals from several locations. In most cases, it accompanied *Sodalis*. Noteworthy, the single characterized specimen from Iran hosted *Pectobacterium* but not *Sodalis*, suggesting potential replacement of the ancestral obligatory nutritional mutualist. The low abundance of *Sodalis* but high of *Pectobacterium* in one of the *P. spumarius* individuals from Kamiennaya Balka, Russia, and one of *P. italosignus* from Sicily might indicate a similar process (Fig. 3A). Similarly, *Sodalis* is absent from a single Greek specimen of *P. signatus*, which in turn hosts abundant *Rickettsia*, an alphaproteobacterium that may contribute to the host nutrition in some cases (Driscoll *et al*., 2017). Interestingly, *Rickettsia* was abundant in almost all populations of *Philaenus* species other than *P. spumarius*, but in this dominant species, only a single individual from Spain was infected. In contrast, *Spiroplasma* was found only in *P. spumarius* and only in the South-Western mitochondrial clade, with high relative abundance (>10%) in the six infected specimens.

**Figure 3.**
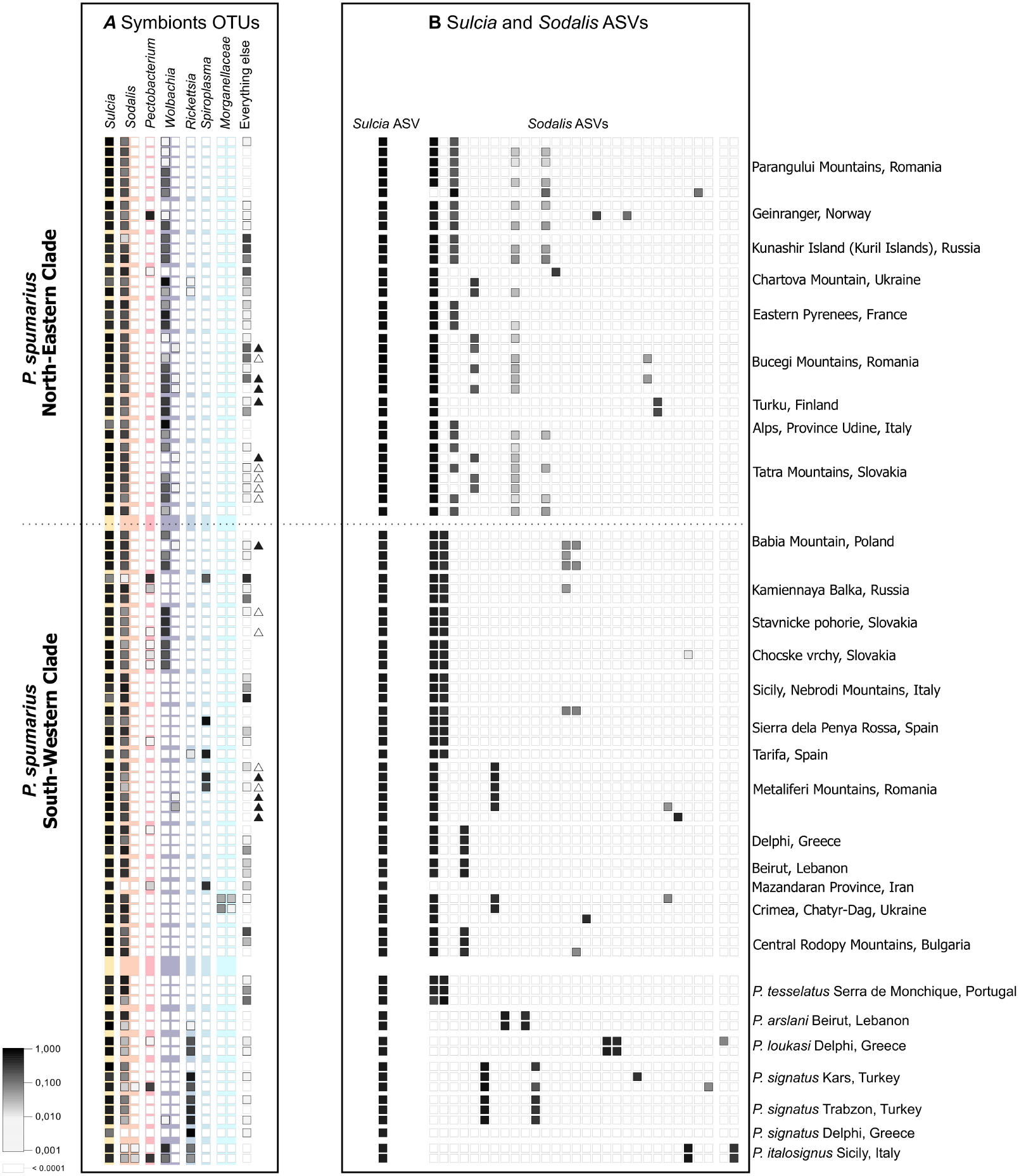
**A** - The distribution of dominant *Philaenus* symbionts across experimental insect populations and species. A. The relative abundance of dominant bacterial OTUs in insect bacterial communities. Triangles next to the heatmap rows indicate the presence of *V. aucta* COI reads in the library: black ones relative abundance >1%, white ones relative abundance <1%. **B** - The relative abundance of dominant ASVs within the two most abundant symbiont OTUs. The shade of grey represents the relative abundance of the given ASV in the *Sulcia* or *Sodalis* OTU, respectively.

In our dataset, we have found two OTUs of *Wolbachia*, one of which was much more prevalent. This dominant OTU showed a specific geographic pattern, where in the North-Eastern clade, it infected 25 out of 35 specimens (65.7%), but in the South-western clade, only 9 examined specimens out of 39 (23.1%) (above the 1% relative abundance threshold). *Wolbachia* infection was significantly more common in the North-Eastern clade (GLM, F1,20.01 = 9.25, p = 0.0064). We also detected *Wolbachia* in one specimen of *P. signatus* and two of *P. italosignus* (Suppl. Table 4). A particularly interesting case was the second *Wolbachia* OTU, much less widespread or abundant when present. All specimens hosting this OTU bore a COI signal of the parasitoid fly *Verralia aucta*, strongly suggesting that this less abundant *Wolbachia* OTU signal is likely of parasitoid origin (Fig. 3A). Specifically, 5 out of 10 parasitoid-positive specimens in the North-Eastern clade and 3 out of 9 in the South-Western clade contained reads of the second *Wolbachia* OTU.

The remaining bacteria in our dataset show less clear patterns. Two OTUs assigned to the family Morganellaceae were present at low abundances in two specimens of *P. spumarius* from Crimea. Surprisingly, we found reads assigned to *Buchnera* in six specimens of *P. spumarius*, verifying the sequence identity to the obligatory nutritional endosymbiont of *Aphis fabae* through BLAST searches against the NCBI nt database. However, with its scattered distribution and the maximum abundance in any library not exceeding 2.5%, we conclude that this is environmental contamination. Other OTUs were generally both relatively rare and low-abundance when present (maximum abundance in any of the libraries <5%) and often assigned to taxa known to be of environmental origins, including *Pseudomonas, Sphingomonas, Staphylococcus*, or members of Rhizobiaceae family. At the same time, we did not detect any known pathogens of plants spread by *Philaenus* spittlebugs such as *Phytoplasma* or *Xylella* (Maejima *et al*., 2014; Sicard *et al*., 2018).

### Genotype-resolution 16S rRNA data highlight host-symbiont interaction dynamics

The single-nucleotide-resolution data for these dominant symbionts revealed that all species and individuals harbor the same ASV of the slow-evolving symbiont *Sulcia* (Fig. 3B). Other symbionts were more variable.

In the case of *Sodalis*, across all individuals, we identified 33 ASVs, of which 12 represented at least 5% of all *Sodalis* reads in at least one sample and were inspected more closely. We found that all specimens contained between 2 and 4 of these more abundant variants of *Sodalis* 16S rRNA gene. Specifically, *P. spumarius* individuals contained a shared abundant ASV, accompanied by others that varied among individuals. Generally, most specimens from the same population hosted identical *Sodalis* ASV combinations, but in many populations, there were individuals with different combinations (Fig. 3). Also, in multiple cases, some individuals hosted additional ASVs besides the combination found in other individuals from the same population. (Fig. 3B).

In specimens of *P. tesselatus*, the Portuguese species unidentifiable from *P. spumarius* at the COI gene, we observed identical *Sodalis* sequence variants as in *P. spumarius* from Spain (Fig. 3B). However, the remaining species were characterized by clearly different *Sodalis* ASV sets than *P. spumarius* and other species. *Sodalis* genomes have been shown to contain multiple rRNA operons that vary in nucleotide different sequences (Koga *et al*., 2013), suggesting that these ASV combinations may often point at different operons in a single genome rather than at different, co-infecting lineages of *Sodalis*.

In the case of *Philaenus-specific Wolbachia* OTU, we observed between 1 and 5 ASVs per individual. The ASV combinations frequently vary among individuals within a population, some ASVs are unique to certain populations, and a single infected *P. signatus* from Turkey harbors a *Wolbachia* genotype absent anywhere else (Fig. 4). These data suggest *Wolbachia* infection dynamics and polymorphism within and between host populations.

**Figure 4.**
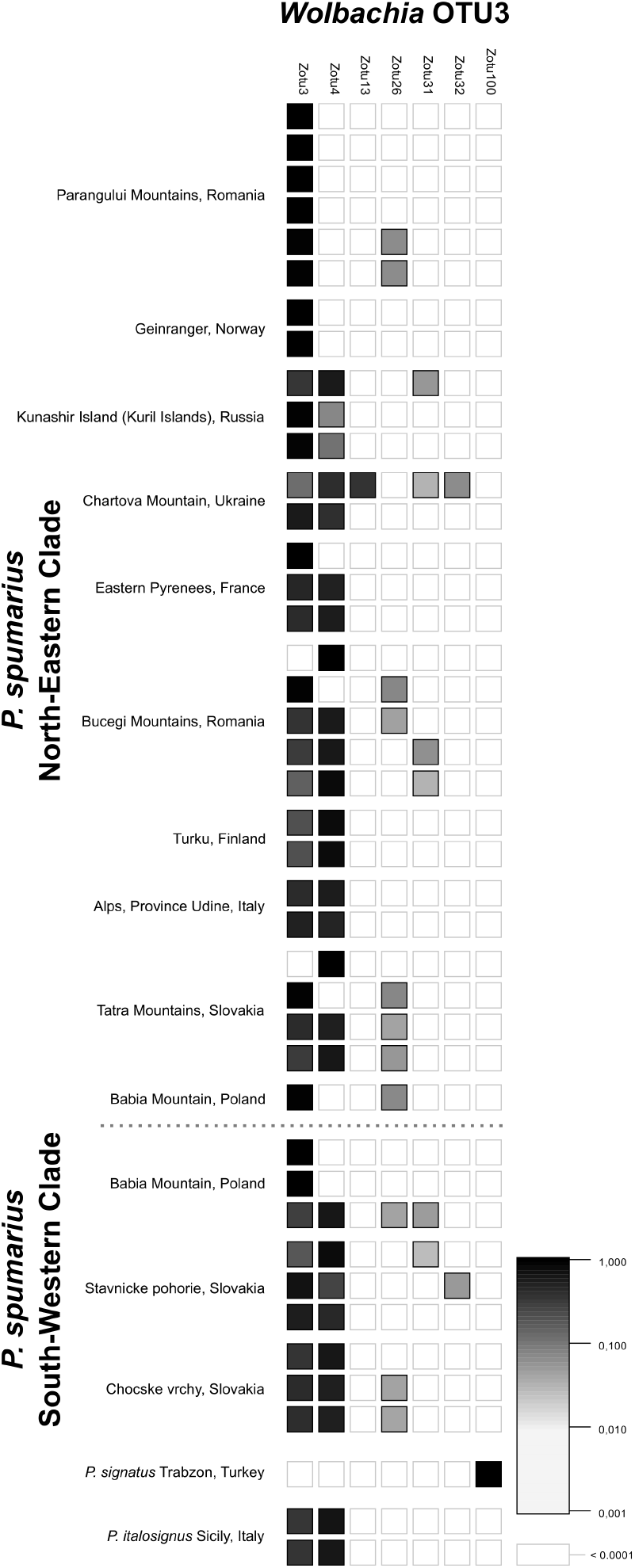
Distribution of *Wolbachia* OTU3 genotypes (ASVs) among *Philaenus* infected specimens.

We also observed variation in ASV-level associations with *Spiroplasma* (Fig. S1) and *Rickettsia* (Fig. S2). In the case of both symbionts, the majority of specimens hosted the same combination of two genotypes (*Spiroplasma*) or a single symbiont genotype (*Rickettsia);* however, for both these microbes, we identified specimens with much higher sequence diversity - four and five ASVs, respectively.

### Obligate symbionts of *P. spumarius* show conserved bacteriocyte-limited location

Histological and ultrastructural observations revealed that obligate symbionts of the spittlebug *P. spumarius* collected from the Polish population reside in the cytoplasm of dedicated insect cells - bacteriocytes (Fig. 5). Bacteriocytes harboring *Sulcia* are mononucleated and create large bacteriomes surrounded by a thick monolayered bacteriome sheath (Fig. 5A-C). The cytoplasm of these bacteriocytes is tightly packed with large, pleomorphic *Sulcia* cells (Fig. 5A-C). Additionally, in the cytoplasm of some *Sulcia* bacteriocytes, we observed tiny bacteria, which may represent *Wolbachia*, common in the northern clade of the species (Fig. 5D). *Sodalis* symbiont cells are spherical and localized in the cytoplasm of separate large bacteriocytes. However, compared to bacteriocytes with *Sulcia*, they are multi-nucleated and do not form compact bacteriomes. Bacteriocytes with *Sodalis* are grouped into larger clusters, but some are separated by fat body cells (Fig. 5E-F). Ultrastructural analyses have shown numerous lamellar structures in the cytoplasm of each bacteriocyte, which we interpret as *Sodalis* cells undergoing degeneration (Fig. 5G). Our observations correspond well with previous microscopy-based studies of *P. spumarius* symbionts conducted by Buchner (1965) and Koga and colleagues (2013). Unfortunately, despite looking at tissues of over 10 individuals, we could not observe additional symbionts in the studied specimens.

**Figure 5.**
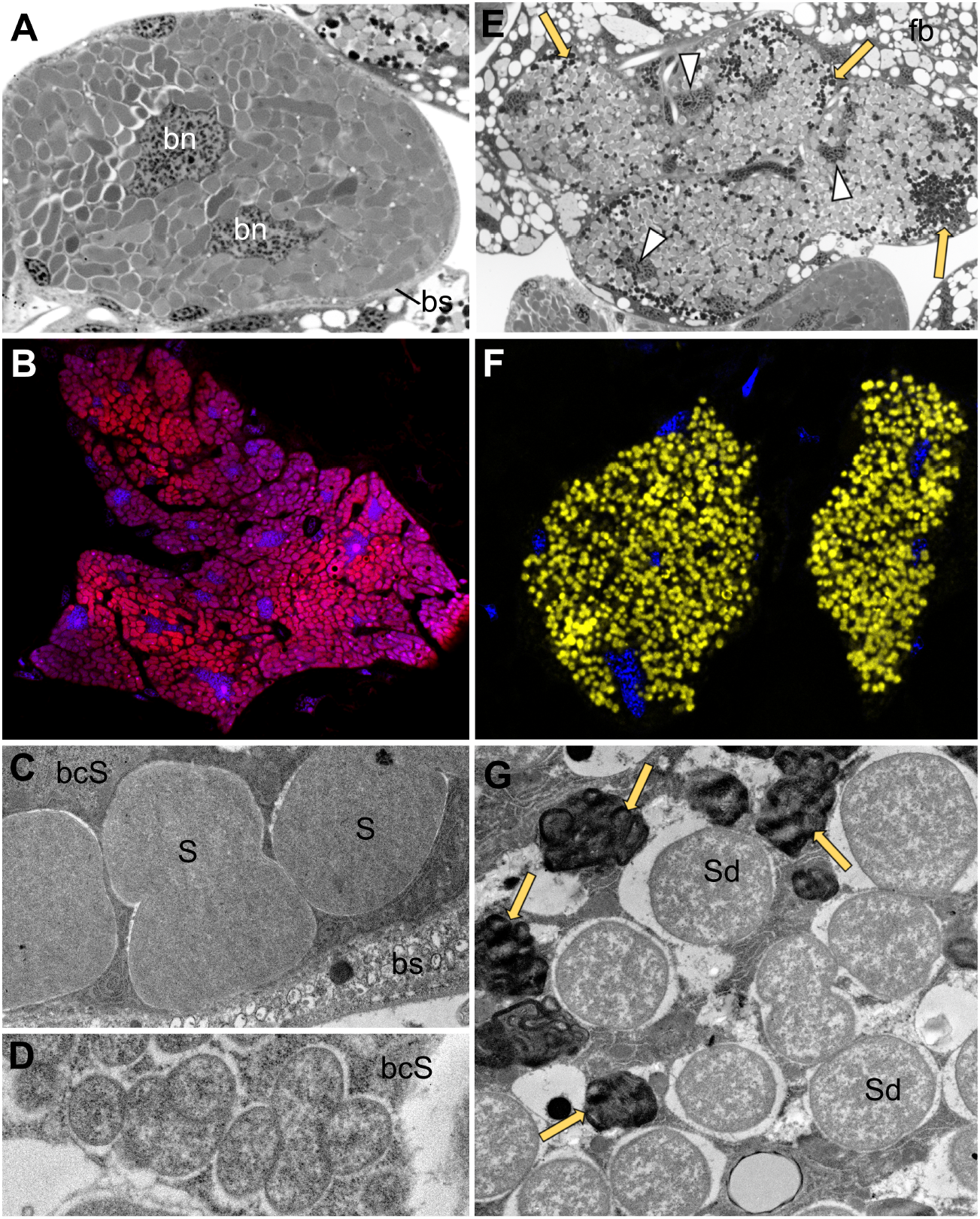
Localization and morphology of symbionts in a population of *P. spumarius* from Kraków, Poland. **A.** Organization of the bacteriome with *Sulcia* symbiont, bn - bacteriocyte nucleus, bs - bacteriome sheath, LM, scale bar = 10 μm. **B.** FISH detection of *Sulcia* symbiont (red); blue represents DAPI staining, FM, scale bar = 10 μm. **C.** Ultrastructure of *Sulcia* (S) cells. bcS - bacteriocyte with *Sulcia* symbiont, bs - bacteriome sheath, TEM, scale bar = 1 μm **D.** Putative *Wolbachia* cells in the cytoplasm of the bacteriocyte harboring *Sulcia* (bcS), TEM, scale bar = 1 μm. **E.** Tissue localization of *Sodalis* symbiont, arrowhead - nucleus, yellow arrow - lamellar structures, scale bar = 10 μm. **F.** *Sodalis* (yellow) visualization using FISH, blue represents DAPI staining, FM, scale bar = 10 μm. **G.** Ultrastructure of *Sodalis* (Sd) symbiont, yellow arrow-degenerated *Sodalis* cells, TEM, scale bar = 1 μm.

## Discussion

### COI amplicon data as the essential framework for the microbiota characterization

The COI amplicon-based characterization of eight morphologically delimited species confirmed the identifications of five (Maryańska-Nadachowska *et al*., 2010). The sixth species, *P. tesselatus*, closely related to *P. spumarius* and distinguishable by the morphology of male genitalia, is known not to be indistinguishable at mitochondrial markers (Maryańska-Nadachowska *et al*., 2012; Rodrigues *et al*., 2014; Seabra *et al*., 2021). The mismatch between morphological and COI-based identification of two other species, both previously found to be clearly distinct at mitochondrial markers from *P. spumarius*, (Maryańska-Nadachowska *et al*., 2010), could point to biological processes such as introgression or laboratory problems such as misidentification or specimen confusion. Nevertheless, their microbial communities are indistinguishable from those of unambiguously identified *P. spumarius*, supporting our decision to reclassify them as this species.

In *P. spumarius*, COI amplicon data provided additional information on the genetic structure below the species level. Our patterns agreed with the prior identifications of two major clades: the South-Western clade ranging from Western Europe, through the Mediterranean, to the Middle East, and the North-Eastern clade ranging from Eastern Asia to Central and Northern Europe (Maryańska-Nadachowska *et al*., 2012; Rodrigues *et al*., 2014; Seabra *et al*., 2021) with the contact zones in the Carpathians, the Alps, the Pyrenees, and the Caucasus (Lis *et al*., 2014; Maryańska-Nadachowskaa *et al*., 2015). These clades have been known to vary in *Wolbachia* infection prevalence (Lis *et al*., 2015), and this study revealed further genetic differences across the sampled geographic regions. Then, even though the 418-bp barcodes provide limited phylogenetic resolution on their own, information on intra-species diversity explains some of the key patterns of symbiont diversity and distribution. We argue that in future microbiome surveys in species of broad interest such as *P. spumarius*, sequencing also host marker genes, combined with genomics datasets for some specimens (Maryańska-Nadachowska *et al*., 2012; Seabra *et al*., 2021), will be essential for the reconstruction of the symbiont distribution, transmission, co-diversification, and replacement patterns, as well as their biological roles. Of particular interest was the signal of the specialized dipteran parasitoid, *V. aucta*, in 24.7% of *P. spumarius* individuals studied. This ratio is comparable to the 17.5% infection prevalence estimates for *P. spumarius* populations from the northern Italy (Molinatto *et al*., 2020), using diagnostic PCR reactions. Then, the COI amplicon sequencing-based screens of individual insects, besides confirming their identity, can provide believable information on some of their biotic associations. Information on the presence and distribution of parasitoid haplotypes within and across *P. spumarius* populations serves as a foundation for a more comprehensive investigation into host-parasitoid-symbiont interaction dynamics.

### Not-so-stable nutritional endosymbioses in *Philaenus* spp

*Sulcia* presence in all studied individuals, and its identity at the targetted rRNA region across all individuals, confirmed its status as a stable nutritional symbiont of spittlebugs (Koga and Moran, 2014). This observation is consistent with its stability across other Cicadomorpha (McCutcheon and Moran, 2010). On the other hand, *Sodalis* relationship with *Philaenus* spittlebugs appears much less stable. It was almost universally prevalent and highly abundant in most individuals, but there were exceptions. Specifically, in two individuals, it seems to have been replaced by other bacteria: *Pectobacterium* or *Rickettsia*. In other individuals, it was present at low abundance, much lower than that of *Pectobacterium, Rickettsia*, or *Wolbachia*, suggestive of complementation or perhaps an ongoing symbiont replacement process. *Symbiopectobacterium* has recently been identified as a versatile microbial clade that has established heritable mutualistic relationships with multiple invertebrate taxa, similar to *Sodalis* (Martinson *et al*., 2020). For example, in parasitic nematode *Howardula aoronymphium*, *Pectobacterium* has been pointed out as a potential nutritional symbiont (Martinson *et al*., 2020). This makes it plausible that in the Iranian *P. spumarius* lineage it has replaced *Sodalis*, taking over its nutritional responsibilities. It would be interesting to investigate whether other *Pectobacterium* infections in our dataset represent early stages of stable associations, cases of vectoring plant pathogens similar to *P. carotovorum* (Mansfield *et al*., 2012) pathogenic infections, or perhaps sporadic, temporary infections with environmentally sourced bacteria.

Interestingly, another case of apparent *Sodalis* loss is a specimen of *P. signatus* from Greece, where *Sulcia* is accompanied by abundant *Rickettsia*. This is surprising as there is no direct evidence that *Rickettsia* can be an obligatory nutritional symbiont in sap-feeding insects, despite the presence of genomic evidence that in some blood-feeders, they may contribute vitamins (Hunter *et al*., 2015). Although there is no particular *Rickettsia* clade correlated with feeding habits, phloem-feeding insects have been considered as an infection hotspot for this microbe (Pilgrim *et al*., 2021).

Recent genomics-enabled data from mealybugs and planthoppers suggest relatively high turnover rates of gammaproteobacterial symbionts such as *Sodalis* during the host evolution (Husnik and McCutcheon, 2016; McCutcheon *et al*., 2019; Michalik *et al*., 2021). Our data suggest the substantial variation among, but also within populations of *P. spumarius*. While 16S rRNA data do not allow us to resolve the phylogenetic relationships among *Sodalis* symbionts from different hosts, especially if only a short gene region is considered, the set of *Sodalis* ASVs, likely representing different operons in the bacterial genome (Koga *et al*., 2013) can apparently serve as a sort of “barcode”, highlighting cases when the symbiosis has likely changed. Specifically, differences in *Sodalis* ASV sets among two insects suggest either divergence in their symbiont genome sequences, infection with independently derived strains, or perhaps different combinations of straisn in case of multiple infections. That way, the variation in ASV sets that we observe among *Philaenus* species, populations, and especially among individuals within a population, indicate either rapid sequence evolution of this symbiont, frequent co-infections with additional strains that result in multiple infections, common symbiont replacements, or a combination of all three processes (McCutcheon *et al*., 2019). Future genomics-enabled work will allow us to distinguish among these scenarios in what appears as a surprisingly dynamic system.

### The diversity and distribution of *Philaenus* facultative endosymbioses

We observed clear patterns in the distribution of dominant facultative endosymbionts, *Wolbachia* (two OTUs), *Rickettsia*, and *Spiroplasma*, across species and populations. Neither symbiont was present in all populations of the studied species, contrary to the results by Kapantaidaki et al. (2021) for Greek populations.

*Wolbachia*, the most broadly distributed insect facultative endosymbiont, is common in *P. spumarius* but its distribution shows clear geographic stratification. The frequencies of *Wolbachia* infection that we estimated in the North-Eastern and South-Western clades of *P. spumarius* are nearly identical to values reported by Lis et al. (2015) on a much larger sampling (c. 70%, and c. 20%, respectively, the latter almost exclusively in the contact zone). This proves that *Wolbachia* diagnostic screens and characterization using Multilocus Sequence Typing (MLST) (Baldo *et al*., 2006) provide concordant results to 16S rRNA amplicon sequencing. At the same time, our simultaneous sequencing of insect COI and bacterial 16S rRNA gene amplicons may explain some of Lis’s and colleagues (2015) observations. We suspect that the previously undescribed *Wolbachia* MLST profiles that they reported in populations in which only single individuals were infected represent *Wolbachia* of parasitoid origin, corresponding to our OTU 25. With parasitoids regarded as a vector for the horizontal transmission of *Wolbachia*, it would be interesting to investigate the nature of this infection further (Ahmed *et al*., 2015). Although Kapantaidaki et al. (2021) reported a high infection rate of *Rickettsia* in populations of *P. spumarius*, we found it rare in this species. However, our findings are congruent in terms of *Rickettsia* that reaches high abundance within infected Philaenus individuals. Further, we may be the first to report *Spiroplasma* infection in the *Philaenus* group. The fourth widespread arthropods reproduction manipulator – *Cardinium*, was absent in our dataset as well as all datasets published previously, adding to evidence on the absence or rarity of this symbiont in *Philaenus* species.

The patterns of the distribution of facultative endosymbionts, especially *Wolbachia* and Rickettsia, may be linked to their fitness effects. Both these symbionts have been linked to their hosts’ thermal susceptibility. According to Hague et al. (2020), some strains of *Wolbachia* supergroup A may shift host preferences toward cooler temperatures, and strains belonging to supergroup B do the opposite. However, those claims do not seem to be broadly supported by data from insects other than *Drosophila* spp. In P. *spumarius*, however, we find the opposite patterns: *Wolbachia* prevalent in the North-Eastern clade (distributed in colder areas) belongs to the supergroup B.

On the other hand, *Rickettsia* may confer protection against heat shock, as demonstrated in whitefly *Bemisia tabaci* (Brumin *et al*., 2011) and, to an extent, in aphids (Montllor *et al*., 2002). Such adaptive properties could explain patterns of infection observed in the *Philaenus* species, namely the presence of *Rickettsia* in specimens from southern regions. However, both these symbionts have multiple other fitness effects that could have contributed to their observed distribution. For example, the combination of reproductive manipulation and positive effects on multiple life-history traits seems to explain the rapid spread of *Wolbachia* across Australian and North American populations of *Drosophila simulans* (Kriesner *et al*., 2013). The same effect may have enabled *Rickettsia* spread in American populations of the whitefly *Bemisia tabaci* (Himler *et al*., 2011). On the other hand, in aphids, *Rickettsia* can protect aphids against pathogenic fungi, perhaps justifying its prevalence in populations when entomopathogenic pressure is high (Łukasik *et al*., 2013a), with its negative effects on fecundity and longevity ameliorated by co-infections with other microbes (Łukasik *et al*., 2013b). Unfortunately, we know little about facultative symbiont roles or fitness effects in Auchenorrhyncha. They are likely to vary substantially among host species and symbiont clades and genotypes, perhaps explaining different *Rickettsia* genotype associations among the North-Eastern and South-Western clades.

Noteworthy, we did not observe the occurrence of some of the facultative endosymbionts described in Greek populations *P. spumarius* described by Kapantaidaki et al. (2021), such as *Arsenophonus* and *Hamiltonella*. Since specimens used in both studies were collected in different sites almost a decade apart, there is a possibility that some of the Greek populations of *P. spumarius* were colonized by the new endosymbionts (Smith *et al*., 2015). Another explanation is that only some populations of *P. spumarius* in Greece are infected by these bacteria, and our sampling (just three individuals) omitted these sites.

On the other hand, since the authors provide information only about a signal and not about the abundance of *Hamiltonella* and *Arsenophonus*, we cannot entirely exclude the environmental origin of that signal. The signal of *Buchnera*, a strictly heritable nutritional endosymbiont of aphids that seems exceedingly unlikely to successfully colonize divergent host such as *P. spumarius* and *P. tesselatus* where we observed it, almost certainly represents some sort of contamination rather than any lasting association. Contamination with *Buchnera-* containing honeydew produced by aphids feeding on the same plant is a possibility, laboratory contamination is another. The knowledge of the biology of diverse insect-associated bacteria helps identify such suspicious cases and is critical for interpreting the data on symbiont distributions.

### Parasitoid effects on host-associated microbial community profiles associations

To our knowledge, we may be the first to show the additional signal of parasitoids in the COI amplicon sequencing data for wild insects and at the same time, its contribution to the overall endosymbiotic signal in microbial community profiles. We obtained information about the parasitoid’s presence, abundance, and mitochondrial genotype alongside information on the mitochondrial diversity of the host. It is clear that molecular methods have opened up new avenues for studying host-parasitoid interactions, with tools such as diagnostic PCR and qPCR assays and sequencing parasitoid marker genes unraveling interaction dynamics and cryptic diversity (Hrček and Godfray, 2015). We argue that COI gene amplicon sequencing shows particular promise as a way of simultaneously obtaining information about the presence of parasitoids on the one hand and their diversity on the other (Šigut *et al*., 2017; Sow *et al*., 2019).

Being able to use a single pair of broad-spectrum primers to simultaneously acquire host genotype information, parasitoid presence, species, and genotype, and host-parasitoid read number ratio as a proxy of parasitoid size does open up exciting avenues for high-throughput study of multitrophic interactions in diverse natural communities. In particular, parasitoid infections may be an important explanatory variable for determining microbial composition, whether by preferential parasitism of hosts that harbor specific combinations of microbes, a change to the host microbiota as a result of parasitism or by parasitoids contributing their microbes to the community profile (Fredensborg *et al*., 2020; Vorburger, 2022). At the same time, symbiotic bacteria are increasingly regarded as shaping host-parasitoid interaction dynamics (Vorburger and Perlman, 2018).

Across these three types of information - host identity confirmation, host genotype information, and the dissection of parasitoid association - COI amplicon data for individual insects are emerging as a groundbreaking tool for the study of microbiota in natural communities.

## Conclusions

The combination of broad specimen sampling with amplicon data for two marker genes proved to be a powerful way of reconstructing host-symbiont relationships, substantially expanding our understanding of a widespread, ecologically and economically significant insect clade.COI amplicon data validated morphology-based species identification and reconstructed parasitoid infections, contributing to the host microbiota signal. OTU-level characterization of microbial communities allowed the reconstruction of broader patterns related to symbiont distribution, whereas genotype-resolution data provided more specific insights into patterns and processes. These complementary approaches have confirmed the stability of the ancient nutritional endosymbiosis with *Sulcia*, the more versatile and dynamic nature of its cosymbiont *Sodalis*, and the geographic and host-phylogenetic structuring of their facultative endosymbioses. Parasitoid infections and likely environmental contaminaton help explain more of the observed variation. And while genomics or experimental approaches are necessary to fully comprehend the biological significance of the observed patterns, the multitarget amplicon sequencing has proven to be a highly effective way or characterizing broader patterns.

## Acknowledgments

The project was supported by the Polish National Agency for Academic Exchange grant PPN/PPO/2018/1/00015, and Polish National Science Centre grant 2018/31/B/NZ8/01158.

## Availability of data

The raw sequence has been deposited in the Sequence Read Archive of the National Centre for Biotechnology Information with the accession number: PRJNA832912

## Supplementary materials

**Supplementary Table 1.**

Collection details.

**Supplementary Table 2.**

Protocols.

**Supplementary Table 3.**

COI OTU table.

**Supplementary Table 4.**

16S OTU table.

**Supplementary Table 5.**

Decontamination statistics table.

**Supplementary Figure 1.**

The distribution of *Spiroplasma* OTU6 genotypes (ASVs).

**Supplementary Figure 2.**

The distribution of *Rickettsia* OTU4 genotypes (ASVs).

## References

Ahmed, M.Z., Li, S.-J., Xue, X., Yin, X.-J., Ren, S.-X., Jiggins, F.M., et al. (2015) The Intracellular Bacterium *Wolbachia* Uses Parasitoid Wasps as Phoretic Vectors for Efficient Horizontal Transmission. PLOS Pathogens 11: e1004672.

Apprill, A., McNally, S., Parsons, R., and Weber, L. (2015) Minor revision to V4 region SSU rRNA 806R gene primer greatly increases detection of SAR11 bacterioplankton. Aquatic Microbial Ecology 75: 129–137.

Baldo, L., Dunning Hotopp, J.C., Jolley, K.A., Bordenstein, S.R., Biber, S.A., Choudhury, R.R., et al. (2006) Multilocus Sequence Typing System for the Endosymbiont *Wolbachia* pipientis. Applied and Environmental Microbiology 72: 7098–7110.

Bennett, G.M. and Moran, N.A. (2015) Heritable symbiosis: the advantages and perils of an evolutionary rabbit hole. Proceedings of the National Academy of Sciences 112: 10169–10176.

Brumin, M., Kontsedalov, S., and Ghanim, M. (2011) *Rickettsia* influences thermotolerance in the whitefly *Bemisia tabaci* B biotype. Insect Science 18: 57–66.

Buchner, P. (1965) Endosymbiosis of animals with plant microorganisms, Interscience Publishers.

Chiel, E., Inbar, M., Mozes-Daube, N., White, J.A., Hunter, M.S., and Zchori-Fein, E. (2009) Assessments of Fitness Effects by the Facultative Symbiont *Rickettsia* in the Sweetpotato Whitefly (Hemiptera: Aleyrodidae). Annals of the Entomological Society of America 102: 413–418.

Douglas, A.E. (2016) How multi-partner endosymbioses function. Nat Rev Microbiol 14: 731–743.

Douglas, A.E. (2009) The microbial dimension in insect nutritional ecology. Functional Ecology 23: 38–47.

Driscoll, T.P., Verhoeve, V.I., Guillotte, M.L., Lehman, S.S., Rennoll, S.A., Beier-Sexton, M., et al. (2017) Wholly *Rickettsia!* Reconstructed Metabolic Profile of the Quintessential Bacterial Parasite of Eukaryotic Cells. mBio 8: e00859–17.

Edgar, R.C. (2016) UNOISE2: improved error-correction for Illumina 16S and ITS amplicon sequencing. bioRxiv 081257.

Elbrecht, V., Braukmann, T.W.A., Ivanova, N.V., Prosser, S.W.J., Hajibabaei, M., Wright, M., et al. (2019) Validation of COI metabarcoding primers for terrestrial arthropods. PeerJ 7: e7745.

Engel, P. and Moran, N.A. (2013) The gut microbiota of insects – diversity in structure and function. FEMS Microbiol Rev 37: 699–735.

Feldhaar, H. (2011) Bacterial symbionts as mediators of ecologically important traits of insect hosts. Ecological Entomology 36: 533–543.

Formisano, G., Iodice, L., Cascone, P., Sacco, A., Quarto, R., Cavalieri, V., et al. (2022) *Wolbachia* infection and genetic diversity of Italian populations of *Philaenus* spumarius, the main vector of *Xylella fastidiosa* in Europe. PLOS ONE 17: e0272028.

Fredensborg, B.L., Fossdal í Kálvalíð, I., Johannesen, T.B., Stensvold, C.R., Nielsen, H.V., and Kapel, C.M. (2020) Parasites modulate the gut-microbiome in insects: A proof-of-concept study. Plos one 15: e0227561.

Gerardo, N.M. and Parker, B.J. (2014) Mechanisms of symbiont-conferred protection against natural enemies: an ecological and evolutionary framework. Current Opinion in Insect Science 4: 8–14.

Glenn, T.C. (2011) Field guide to next-generation DNA sequencers. Molecular Ecology Resources 11: 759–769.

Godefroid, M., Morente, M., Schartel, T., Cornara, D., Purcell, A., Gallego, D., et al. (2021) Climate tolerances of *Philaenus spumarius* should be considered in risk assessment of disease outbreaks related to Xylella fastidiosa. J Pest Sci.

Hague, M.T.J., Caldwell, C.N., and Cooper, B.S. (2020) Pervasive Effects of *Wolbachia* on Host Temperature Preference. mBio 11:.

Hedges, L.M., Brownlie, J.C., O’Neill, S.L., and Johnson, K.N. (2008) *Wolbachia* and Virus Protection in Insects. Science 322: 702–702.

Himler, A.G., Adachi-Hagimori, T., Bergen, J.E., Kozuch, A., Kelly, S.E., Tabashnik, B.E., et al. (2011) Rapid spread of a bacterial symbiont in an invasive whitefly is driven by fitness benefits and female bias. science 332: 254–256.

Hosokawa, T., Koga, R., Kikuchi, Y., Meng, X.-Y., and Fukatsu, T. (2010) *Wolbachia* as a bacteriocyte-associated nutritional mutualist. PNAS 107: 769–774.

Hrček, J. and Godfray, H.C.J. (2015) What do molecular methods bring to host-parasitoid food webs? Trends Parasitol 31: 30–35.

Hunter, D.J., Torkelson, J.L., Bodnar, J., Mortazavi, B., Laurent, T., Deason, J., et al. (2015) The *Rickettsia* Endosymbiont of *Ixodes pacificus* Contains All the Genes of De Novo Folate Biosynthesis. PLOS ONE 10: e0144552.

Hurst, G.D. and Jiggins, F.M. (2000) Male-killing bacteria in insects: mechanisms, incidence, and implications. Emerg Infect Dis 6: 329–336.

Husnik, F. and McCutcheon, J.P. (2016) Repeated replacement of an intrabacterial symbiont in the tripartite nested mealybug symbiosis. PNAS 113: E5416–E5424.

Jaenike, J., Unckless, R., Cockburn, S.N., Boelio, L.M., and Perlman, S.J. (2010) Adaptation via Symbiosis: Recent Spread of a *Drosophila* Defensive Symbiont. Science 329: 212–215.

Ju, J.-F., Bing, X.-L., Zhao, D.-S., Guo, Y., Xi, Z., Hoffmann, A.A., et al. (2020) *Wolbachia* supplement biotin and riboflavin to enhance reproduction in planthoppers. ISME J 14: 676–687.

Kambris, Z., Blagborough, A.M., Pinto, S.B., Blagrove, M.S.C., Godfray, H.C.J., Sinden, R.E., and Sinkins, S.P. (2010) *Wolbachia* Stimulates Immune Gene Expression and Inhibits *Plasmodium* Development in *Anopheles gambiae*. PLOS Pathogens 6: e1001143.

Kapantaidaki, D.E., Antonatos, S., Evangelou, V., Papachristos, D.P., and Milonas, P. (2021) Genetic and endosymbiotic diversity of Greek populations of *Philaenus spumarius*, *Philaenus signatus* and *Neophilaenus campestris*, vectors of *Xylella fastidiosa*. Sci Rep 11: 3752.

Knight, R., Vrbanac, A., Taylor, B.C., Aksenov, A., Callewaert, C., Debelius, J., et al. (2018) Best practices for analysing microbiomes. Nature Reviews Microbiology 16: 410–422.

Koga, R., Bennett, G.M., Cryan, J.R., and Moran, N.A. (2013) Evolutionary replacement of obligate symbionts in an ancient and diverse insect lineage. Environmental Microbiology 15: 2073–2081.

Koga, R. and Moran, N.A. (2014) Swapping symbionts in spittlebugs: evolutionary replacement of a reduced genome symbiont. ISME J 8: 1237–1246.

Kriesner, P., Hoffmann, A.A., Lee, S.F., Turelli, M., and Weeks, A.R. (2013) Rapid sequential spread of two Wolbachia variants in Drosophila simulans. PLoS Pathog 9:.

Leigh, J.W. and Bryant, D. (2015) POPART: full-feature software for haplotype network construction. Methods in Ecology and Evolution 6: 1110–1116.

Lis, A., Maryańska-Nadachowska, A., and Kajtoch, Ł. (2015) Relations of *Wolbachia* Infection with Phylogeography of *Philaenus spumarius* (Hemiptera: Aphrophoridae) Populations Within and Beyond the Carpathian Contact Zone. Microb Ecol 70: 509–521.

Lis, A., Maryańska-Nadachowska, A., Lachowska-Cierlik, D., and Kajtoch, Ł. (2014) The Secondary Contact Zone of Phylogenetic Lineages of the *Philaenus spumarius* (Hemiptera: Aphrophoridae): An Example of Incomplete Allopatric Speciation. Journal of Insect Science 14:.

Liu, X.-D. and Guo, H.-F. (2019) Importance of endosymbionts *Wolbachia* and *Rickettsia* in insect resistance development. Current Opinion in Insect Science 33: 84–90.

Łukasik, Piotr, Asch, M. van, Guo, H., Ferrari, J., and Godfray, H.C.J. (2013) Unrelated facultative endosymbionts protect aphids against a fungal pathogen. Ecology Letters 16: 214–218.

Łukasik, P., Guo, H., Asch, M. van, Ferrari, J., and Godfray, H.C.J. (2013) Protection against a fungal pathogen conferred by the aphid facultative endosymbionts *Rickettsia* and *Spiroplasma* is expressed in multiple host genotypes and species and is not influenced by co-infection with another symbiont. Journal of Evolutionary Biology 26: 2654–2661.

Luo, J., Cheng, Y., Guo, L., Wang, A., Lu, M., and Xu, L. (2021) Variation of gut microbiota caused by an imbalance diet is detrimental to bugs’ survival. Science of The Total Environment 771: 144880.

Maejima, K., Oshima, K., and Namba, S. (2014) Exploring the phytoplasmas, plant pathogenic bacteria. J Gen Plant Pathol 80: 210–221.

Mansfield, J., Genin, S., Magori, S., Citovsky, V., Sriariyanum, M., Ronald, P., et al. (2012) Top 10 plant pathogenic bacteria in molecular plant pathology. Molecular Plant Pathology 13: 614–629.

Martinson, V.G., Gawryluk, R.M.R., Gowen, B.E., Curtis, C.I., Jaenike, J., and Perlman, S.J. (2020) Multiple origins of obligate nematode and insect symbionts by a clade of bacteria closely related to plant pathogens. PNAS 117: 31979–31986.

Maryańska-Nadachowska, A., Drosopoulos, S., Lachowska, D., Kajtoch, Ł., and Kuznetsova, V.G. (2010) Molecular phylogeny of the Mediterranean species of *Philaenus* (Hemiptera: Auchenorrhyncha: Aphrophoridae) using mitochondrial and nuclear DNA sequences. Systematic Entomology 35: 318–328.

Maryańska-Nadachowska, A., Kajtoch, Ł., and Lachowska, D. (2012) Genetic diversity of Philaenus spumarius and P. tesselatus (Hemiptera, Aphrophoridae): implications for evolution and taxonomy. Systematic Entomology 37: 55–64.

Maryańska-Nadachowska, A., Sanaieb, E., and Kajtocha, Ł. (2015) High genetic diversity in southwest Asian populations of *Philaenus spumarius* (Hemiptera: Auchenorrhyncha). Zoology in the Middle East 61: 264–272.

Matsuura, Y., Moriyama, M., Łukasik, P., Vanderpool, D., Tanahashi, M., Meng, X.-Y., et al. (2018) Recurrent symbiont recruitment from fungal parasites in cicadas. PNAS 115: E5970–E5979.

McCutcheon, J.P., Boyd, B.M., and Dale, C. (2019) The Life of an Insect Endosymbiont from the Cradle to the Grave. Current Biology 29: R485–R495.

McCutcheon, J.P., McDonald, B.R., and Moran, N.A. (2009) Convergent evolution of metabolic roles in bacterial co-symbionts of insects. PNAS 106: 15394–15399.

McCutcheon, J.P. and Moran, N.A. (2010) Functional convergence in reduced genomes of bacterial symbionts spanning 200 My of evolution. Genome biology and evolution 2: 708–718.

Michalik, A., Castillo Franco, D., Kobiałka, M., Szklarzewicz, T., Stroiński, A., and Łukasik, P. (2021) Alternative Transmission Patterns in Independently Acquired Nutritional Cosymbionts of Dictyopharidae Planthoppers. mBio 12: e01228–21.

Molinatto, G., Demichelis, S., Bodino, N., Giorgini, M., Mori, N., and Bosco, D. (2020) Biology and Prevalence in Northern Italy of *Verrallia aucta* (Diptera, Pipunculidae), a Parasitoid of *Philaenus spumarius* (Hemiptera, Aphrophoridae), the Main Vector of *Xylella fastidiosa* in Europe. Insects 11: 607.

Montllor, C.B., Maxmen, A., and Purcell, A.H. (2002) Facultative bacterial endosymbionts benefit pea aphids *Acyrthosiphon pisum* under heat stress. Ecological Entomology 27: 189–195.

Moran, N.A., McCutcheon, J.P., and Nakabachi, A. (2008) Genomics and evolution of heritable bacterial symbionts. Annual Review of Genetics 42: 165–190.

Parada, A.E., Needham, D.M., and Fuhrman, J.A. (2016) Every base matters: assessing small subunit rRNA primers for marine microbiomes with mock communities, time series and global field samples. Environmental Microbiology 18: 1403–1414.

Perreau, J. and Moran, N.A. (2021) Genetic innovations in animal–microbe symbioses. Nat Rev Genet 1–17.

Pilgrim, J., Thongprem, P., Davison, H.R., Siozios, S., Baylis, M., Zakharov, E.V., et al. (2021) Torix *Rickettsia* are widespread in arthropods and reflect a neglected symbiosis. GigaScience 10: giab021.

Prodan, A., Tremaroli, V., Brolin, H., Zwinderman, A.H., Nieuwdorp, M., and Levin, E. (2020) Comparing bioinformatic pipelines for microbial 16S rRNA amplicon sequencing. PLOS ONE 15: e0227434.

Rodrigues, A.S.B., Silva, S.E., Marabuto, E., Silva, D.N., Wilson, M.R., Thompson, V., et al. (2014) New Mitochondrial and Nuclear Evidences Support Recent Demographic Expansion and an Atypical Phylogeographic Pattern in the Spittlebug Philaenus spumarius (Hemiptera, Aphrophoridae). PLOS ONE 9: e98375.

Rognes, T., Flouri, T., Nichols, B., Quince, C., and Mahé, F. (2016) VSEARCH: a versatile open source tool for metagenomics. PeerJ 4: e2584.

Salter, S.J., Cox, M.J., Turek, E.M., Calus, S.T., Cookson, W.O., Moffatt, M.F., et al. (2014) Reagent and laboratory contamination can critically impact sequence-based microbiome analyses. BMC Biology 12: 87.

Saponari, M., Loconsole, G., Cornara, D., Yokomi, R.K., De Stradis, A., Boscia, D., et al. (2014) Infectivity and Transmission of *Xylella fastidiosa* by *Philaenus spumarius* (Hemiptera: Aphrophoridae) in Apulia, Italy. Journal of Economic Entomology 107: 1316–1319.

Seabra, S.G., Rodrigues, A.S.B., Silva, S.E., Neto, A.C., Pina-Martins, F., Marabuto, E., et al. (2021) Population structure, adaptation and divergence of the meadow spittlebug, *Philaenus spumarius* (Hemiptera, Aphrophoridae), revealed by genomic and morphological data. PeerJ 9: e11425.

Sicard, A., Zeilinger, A.R., Vanhove, M., Schartel, T.E., Beal, D.J., Daugherty, M.P., and Almeida, R.P.P. (2018) *Xylella fastidiosa:* Insights into an Emerging Plant Pathogen, Annual Reviews.

Šigut, M., Kostovčík, M., Šigutová, H., Hulcr, J., Drozd, P., and Hrček, J. (2017) Performance of DNA metabarcoding, standard barcoding, and morphological approach in the identification of host–parasitoid interactions. PLOS ONE 12: e0187803.

Smith, A.H., Łukasik, P., O’Connor, M.P., Lee, A., Mayo, G., Drott, M.T., et al. (2015) Patterns, causes and consequences of defensive microbiome dynamics across multiple scales. Molecular Ecology 24: 1135–1149.

Smith, A.H., O’Connor, M.P., Deal, B., Kotzer, C., Lee, A., Wagner, B., et al. (2021) Does getting defensive get you anywhere?—Seasonal balancing selection, temperature, and parasitoids shape real-world, protective endosymbiont dynamics in the pea aphid. Molecular Ecology 30: 2449–2472.

Song, H., Buhay, J.E., Whiting, M.F., and Crandall, K.A. (2008) Many species in one: DNA barcoding overestimates the number of species when nuclear mitochondrial pseudogenes are coamplified. Proceedings of the National Academy of Sciences 105: 13486–13491.

Sow, A., Brévault, T., Benoit, L., Chapuis, M.-P., Galan, M., Coeur d’acier, A., et al. (2019) Deciphering host-parasitoid interactions and parasitism rates of crop pests using DNA metabarcoding. Sci Rep 9: 3646.

Truitt, A.M., Kapun, M., Kaur, R., and Miller, W.J. (2019) *Wolbachia* modifies thermal preference in Drosophila melanogaster. Environmental microbiology 21: 3259–3268.

Turelli, M., Cooper, B.S., Richardson, K.M., Ginsberg, P.S., Peckenpaugh, B., Antelope, C.X., et al. (2018) Rapid Global Spread of wRi-like *Wolbachia* across Multiple Drosophila. Curr Biol 28: 963–971.e8.

Vorburger, C. (2022) Defensive Symbionts and the Evolution of Parasitoid Host Specialization. Annual Review of Entomology 67: 329–346.

Vorburger, C. and Perlman, S.J. (2018) The role of defensive symbionts in host–parasite coevolution. Biological Reviews 93: 1747–1764.

Xie, J., Butler, S., Sanchez, G., and Mateos, M. (2014) Male killing *Spiroplasma* protects *Drosophila melanogaster* against two parasitoid wasps. Heredity 112: 399–408.

Zhang, J., Kobert, K., Flouri, T., and Stamatakis, A. (2014) PEAR: a fast and accurate Illumina Paired-End reAd mergeR. Bioinformatics 30: 614–620.

Zug, R. and Hammerstein, P. (2012) Still a Host of Hosts for Wolbachia: Analysis of Recent Data Suggests That 40% of Terrestrial Arthropod Species Are Infected. PLOS ONE 7: e38544.

