## Supplementary figures and images for "Till evolution do us part: The diversity of symbiotic associations across populations of *Philaenus* spittlebugs"

### Supplementary Figure 1

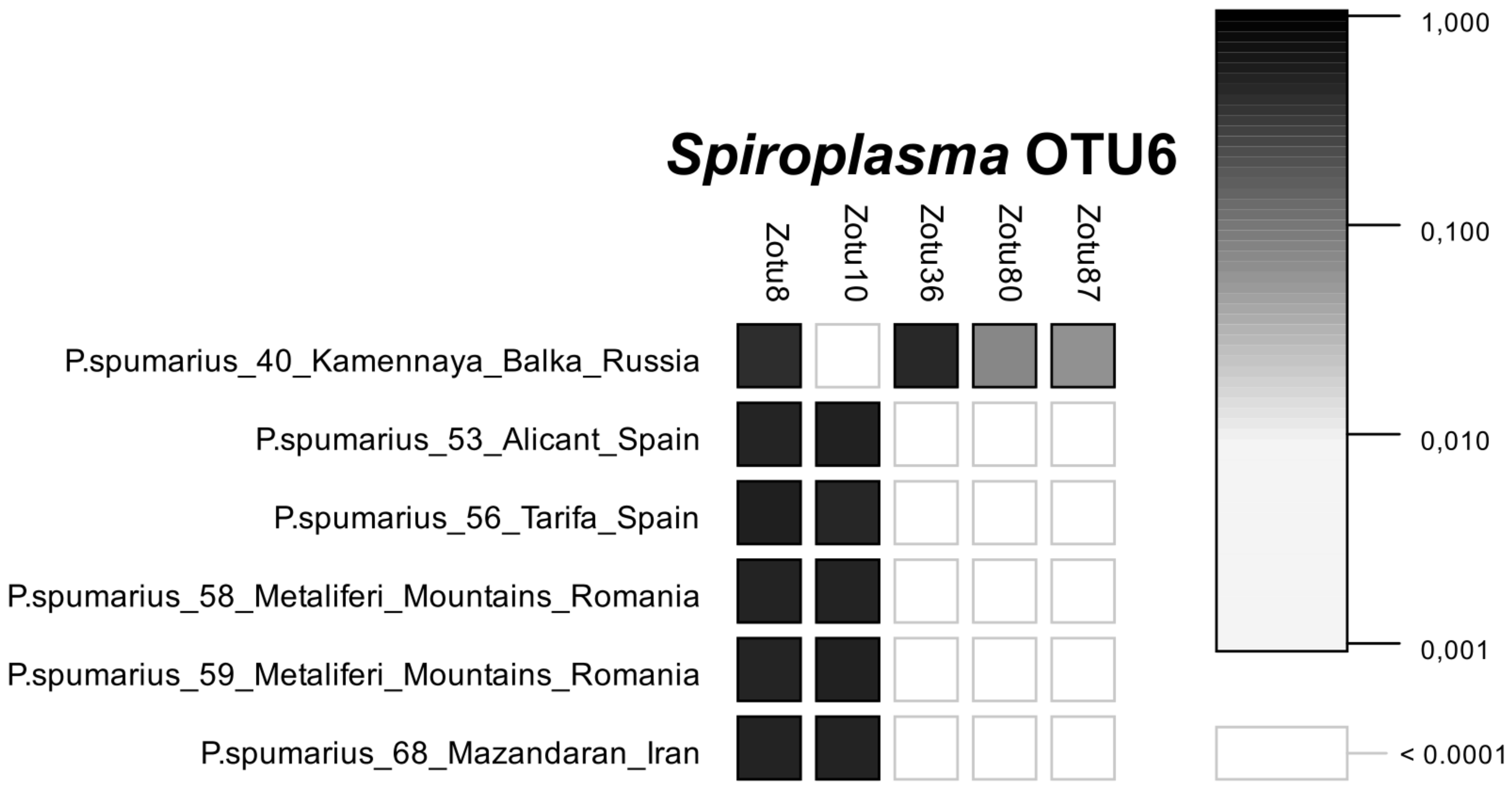

### Supplementary Figure 2

#### *Rickettsia* OTU4

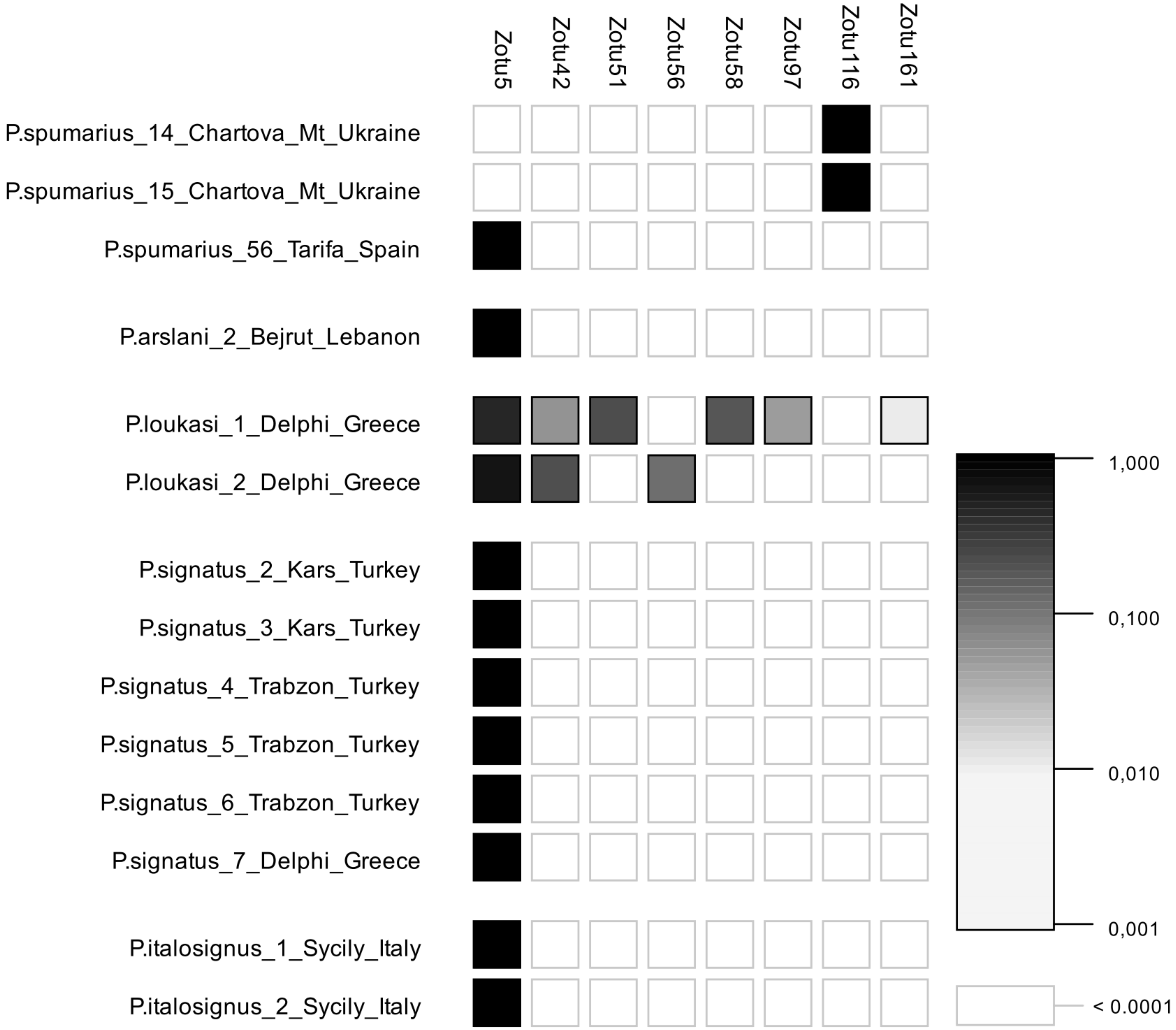
